# The lipid environment determines the activity of the *E. coli* ammonium transporter, AmtB

**DOI:** 10.1101/303065

**Authors:** Gaëtan Dias Mirandela, Giulia Tamburrino, Paul A. Hoskisson, Ulrich Zachariae, Arnaud Javelle

## Abstract

The movement of ammonium across biological membranes is a fundamental process in all living organisms and is mediated by the ubiquitous Amt/Mep/Rh family of transporters. Recent structural analysis and coupled mass spectrometry studies have shown that the *Escherichia coli* ammonium transporter, AmtB, specifically binds phosphatidylglycerol (PG). Upon PG binding, several residues of AmtB undergo a small conformational change, which stabilizes the protein against unfolding. However, no studies have so far been conducted to explore if PG binding to AmtB has functional consequences. Here, we used an in *vitro* experimental assay with purified components together with molecular dynamics simulations to characterise the relation between PG binding and AmtB activity. Firstly, our results indicate that the function of Amt in archaebacteria and eubacteria may differ. Secondly, we show that PG is an essential cofactor for AmtB activity and that in the absence of PG AmtB cannot complete the full translocation cycle. Furthermore, our simulations reveal previously undiscovered PG binding sites on the intracellular side of the lipid bilayer between the AmtB subunits. The possible molecular mechanisms explaining the functional role of PG are discussed.

## Introduction

Ammonium is a vital source of nitrogen for bacteria, fungi and plants and a toxic metabolic waste product for animals (1). Hence, ammonium transport across biological membranes is a process of fundamental importance in all living organisms. In 1994, the first genes encoding ammonium transporters were identified in *Saccharomyces cerevisiae* (*mep* for methylammonium permease) (2) and *Arabidopsis thaliana* (*amt* for ammonium transporter) (3). Later it was shown that the rhesus protein (Rh) is an orthologue of Amt in vertebrates (4) and remarkably, that yeast *mep* mutants can be complemented with the human Rh glycoprotein (5), demonstrating that the Rh protein is a functional ammonium transporter. Following these seminal findings, members of the Amt/Mep/Rh protein family have been identified in almost all sequenced organisms, forming a unique and highly specific family of ammonium transporters (6, 7).

The functional context of Amt/Mep and Rh transporters is highly diverse: bacteria, fungi and plants use Amt/Mep proteins to scavenge ammonium from their environments for biosynthetic assimilation, whereas mammals use the Rh proteins for ammonium detoxification in erythrocytes, kidney and liver tissues (1, 8, 9). Hence, members of the Amt/Mep/Rh family of proteins are associated with various fundamental biological processes. In fungi the dimorphic transition from yeast to filamentation is often related to the virulence of pathogenic species such as *Candida albicans* (10), *Histoplasma capsulatum* (11) and *Cryptococcus neoformans* (12). Fungi possess multiple Mep proteins and it has been shown that in *S. cerevisiae* (13), the plant pathogens *Ustilago maydis* (14, 15), *Fusarium fujikuroi* (16) and the human pathogen *C. albicans* (17) the Mep2 transporters play a key role in the switch to filamentous growth.

In humans, Rh mutations are associated with numerous pathologies. RhAG mutations in red blood cells have been linked to recessive rhesus protein deficiency (18) and Overhydrated Hereditary Stomatocytosis (19) - a rare, dominant inherited haemolytic anaemia. In mouse kidneys, RhCG mutations impair ammonium homeostasis and are associated with distal Renal Tubular Acidosis and male infertility (20). Finally, RhCG has been identified as a candidate gene for early-onset major depressive disorder (21).

The *Escherichia coli* ammonium transporter AmtB is the most widely studied model system to investigate ammonium uptake in the ubiquitous Amt/Mep/Rh protein family (22). AmtB is structurally well characterised, with more than 20 high resolution structures reported in the Protein Data Bank (PDB) to date. Despite this wealth of structural information, the ammonium transport mechanism has not yet been unravelled from these crystal structures, since all the structures show a very similar conformation reflecting the inward-facing state of the protein, irrespective of the presence or absence of ammonium. Recently, mass spectrometric analysis coupled with structural studies defined two specific binding sites for the phosphatidylglycerol (PG) head group in AmtB, which increase protein stability (23). The X-ray structure of AmtB with bound PG reveals distinct conformational changes which reposition some of the protein residues that interact with lipids. More recently, it has been shown that PG can allosterically regulate the interaction between AmtB and the signal transduction protein GlnK (24). Despite these findings, a direct functional role for PG on the transport of ammonium by AmtB has remained unclear.

Here, we couple an *in vitro* assay, based on protein reconstitution in liposomes and surface supported membrane electrophysiology (SSME) measurements with molecular dynamics simulations, to illuminate the effect of lipid composition on AmtB activity. Firstly, our results indicate that the function of Amt in archaebacterial and eubacteria differs. Secondly, we show that PG is an essential cofactor for AmtB activity and that in the absence of PG AmtB cannot complete the full translocation cycle. To our best knowledge, this is the first report highlighting the functional importance of specific lipids for AmtB activity and demonstrating that the high AmtB selectivity for PG lipids is not only important for protein stability but also for the translocation cycle.

### Characterization of AmtB activity by SSME

To measure ammonium transport activity using SSME, we purified AmtB as previously described (25). The protein was incorporated into liposomes containing a mixture of *E. coli* polar lipids/PC at a weight ratio of 2:1. AmtB was reconstituted at a lipid-to-protein ratio (LPR) of 10 (wt/wt). Dynamic light scattering (DLS) analysis confirmed that liposomes and proteoliposomes follow a unimodal size distribution with a mean size of 110 nm (Figure 1A). To examine the orientation of AmtB inserted in the liposomes, we analysed the proteoliposome by Immobilised Metal Affinity Chromatography IMAC. If AmtB is inserted in a right-side-out (RSO) orientation, the C-terminal affinity tag should not be accessible, hence the proteoliposomes should flow through the IMAC matrix. By contrast, if the insertion is orientated inside-out (IO), then the His-tag should be accessible and the proteoliposomes is expected to bind the matrix and thus be present in the elution fraction. As a control, we treated the proteoliposomes in parallel with DDM to solubilise AmtB and analyse it by IMAC using identical conditions as for the analysis of the proteoliposomes. Our analysis showed that about half of the proteoliposomes are present in the flow through and the other half in the elution fraction, demonstrating that AmtB is orientated approximately 50/50% RSO/IO in the proteoliposomes (Figure 1B).

**Figure 1:**
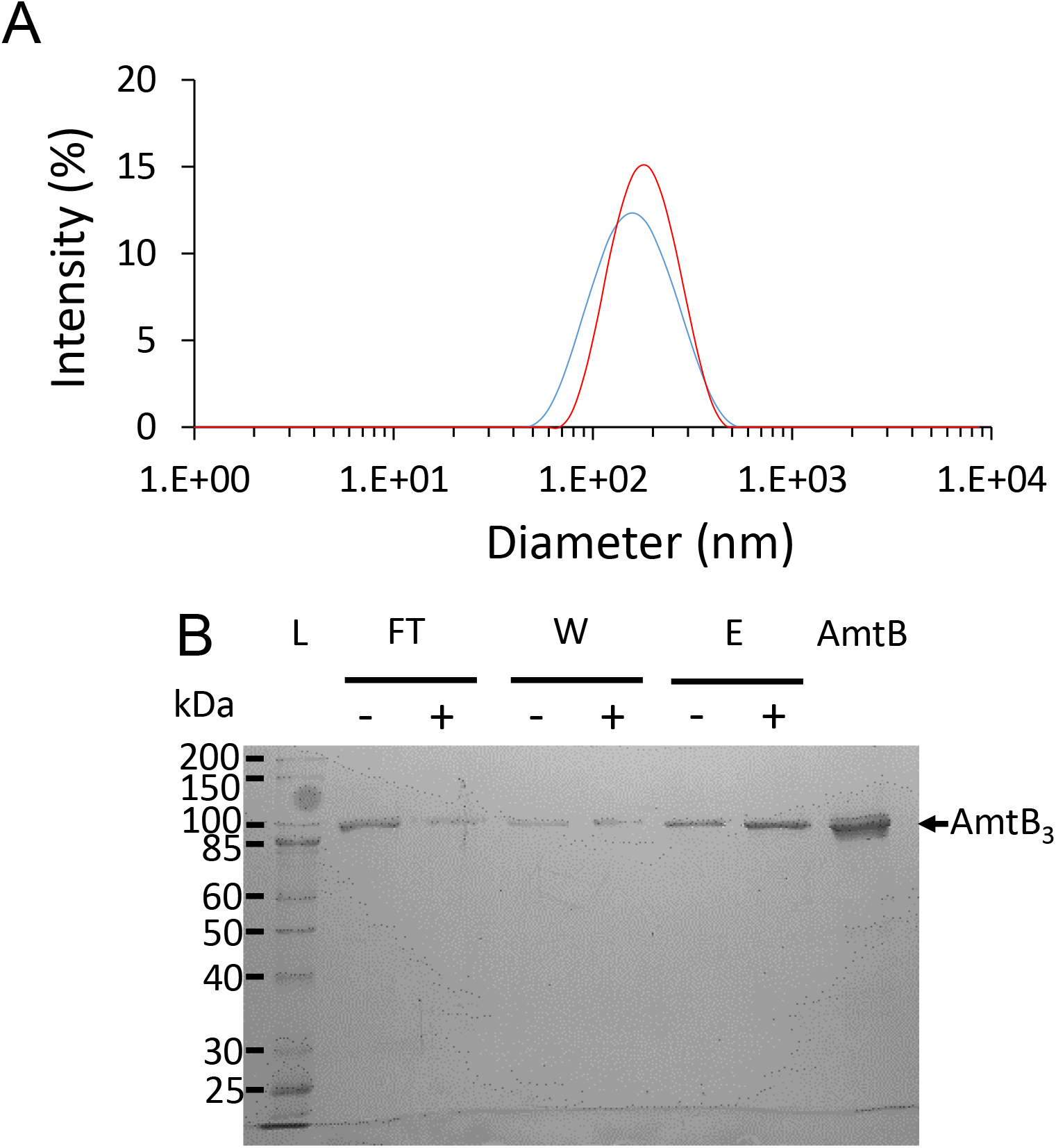
AmtB purification and reconstitution into liposomes. **(A)** DLS analysis of the empty liposomes (blue) and proteo-liposomes (red). **(B)** SDS-PAGE Coomassie Blue-stained gel of the liposomes purified by IMAC after DDM treatment (+) or in absence of DDM (-). FT; flow through, W; wash, E; elution fraction, AmtB; 5mg of pure AmtB used for the reconstitution in the proteoliposomes

SSME analysis after an ammonium pulse of 100 mM revealed a fast positive transient current of 3.3 nA in proteoliposomes, whereas no current was recorded for protein-free liposomes (Figure 2). We show a representative trace in Figure 2, however it should be noted that the amplitude of the transient current differs from sensor to sensor because the number of proteoliposomes coated on the Surface Supported Membrane (SSM) varies (26). In our standard experimental set-up, we measured the current two times on six sensors produced from two independent protein preparations. The average transient current peak measured for a pulse of 100 mM at LPR10 was 3.37±0.26 nA. SSME records transient currents as the charge displacement caused by the translocation of ammonium inside the proteoliposomes creates an outwardly directed negative membrane potential that progressively inhibits the transport cycle. This fast transient current measures both pre-steady-state charge displacement (corresponding to the interaction of ammonium with AmtB) and steady-state charge displacement (describing the continuous turnover during the complete transport cycle of AmtB) (26). To further confirm that the transient currents correspond to the translocation of ammonium into the proteoliposomes rather than a simple interaction between the substrate and the transporters, we investigated the effect of varying the number of transporters per proteoliposome on the transient current. It is expected that the decay time is prolonged with increasing protein in the liposomes if the current represents a complete transport cycle, while it should be independent of the number of transporters per liposome if the current reflects a simple binding interaction between the substrate and the protein (27). To test this, we reconstituted AmtB into liposomes at LPR values of 50, 10 and 5 (wt/wt). After a pulse of 100 mM ammonium, the maximum amplitude of the transient current between LPR 50, 10 and 5 increased from 0.47±0.02 nA to 3.37±0.26 and 7.90±0.35, respectively. More importantly still, the decay rate constant of the second phase increases from 9.5±0.7 s^-1^ to 13.4±1.6 s^-1^ and 18.7±1 s^-1^ (Figure 2). Taken together, these results show that a charge displacement specific to AmtB can be detected and that the current describes the continous turnover of the complete transport cycle. To determine the transport kinetics, the transient currents were measured in proteoliposomes reconstituted at LPR10, following ammonium pulses ranging from 0.024 mM to 100 mM. The peak currents saturated between ammonium pulses of 25-50 mM, hence we normalised our recordings against the current measured at 100 mM. The data were then fitted according to the Michaelis-Menten equation and a *K*_m_ of 0.8±0.1 mM was calculated (r^2^=0.99) (Figure 3B).

**Figure 2:**
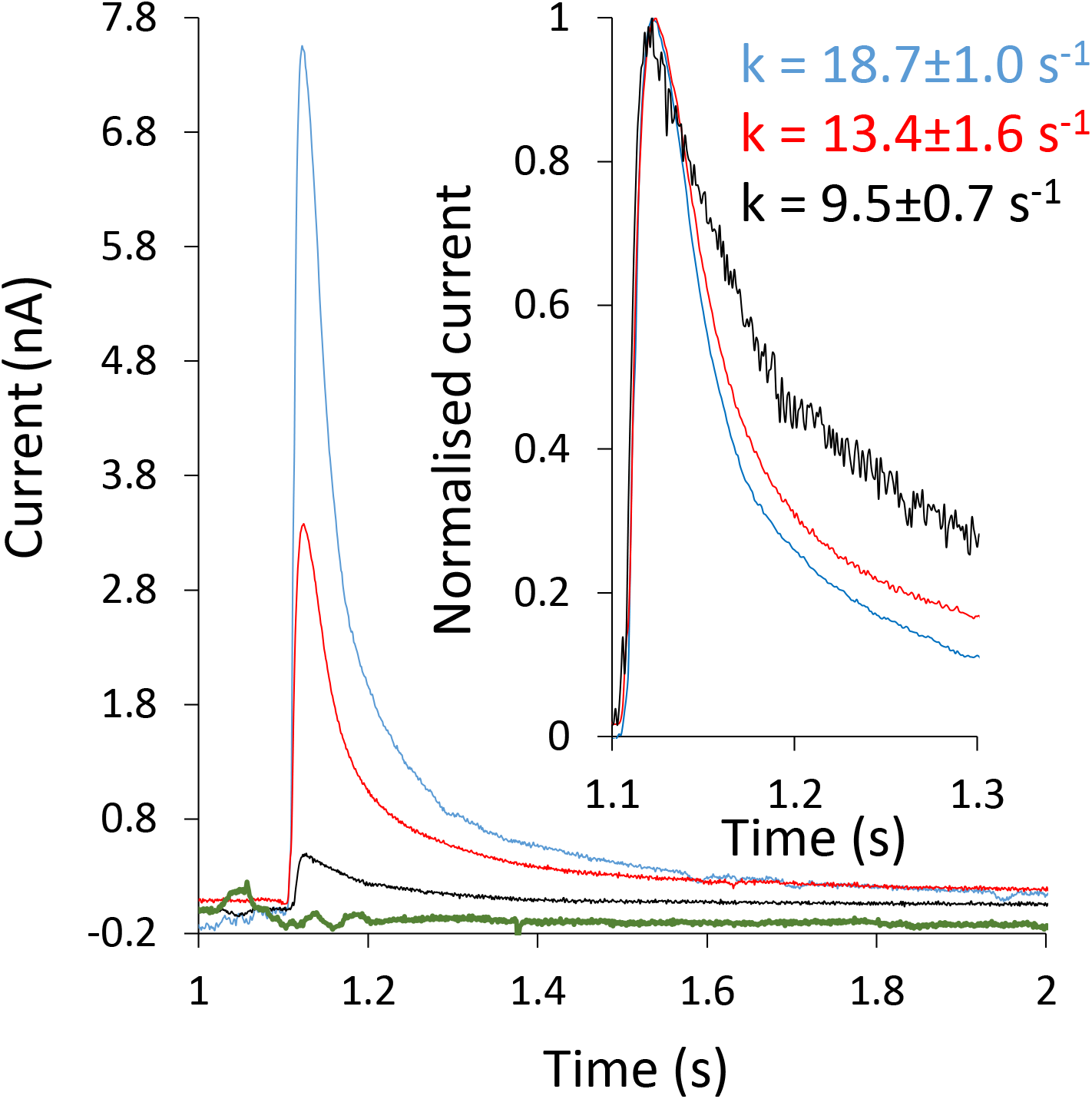
Characterization of AmtB activity. Transient current measured after a 100 mM ammonium jump in empty liposomes (green) or proteoliposomes containing AmtB at a LPR of 50 (black), 10 (red) or 5 (blue). ***Insert***: Normalised current measured in proteoliposomes containing AmtB at a LPR of 50 (black), 10 (red) or 5 (blue).

**Figure 3:**
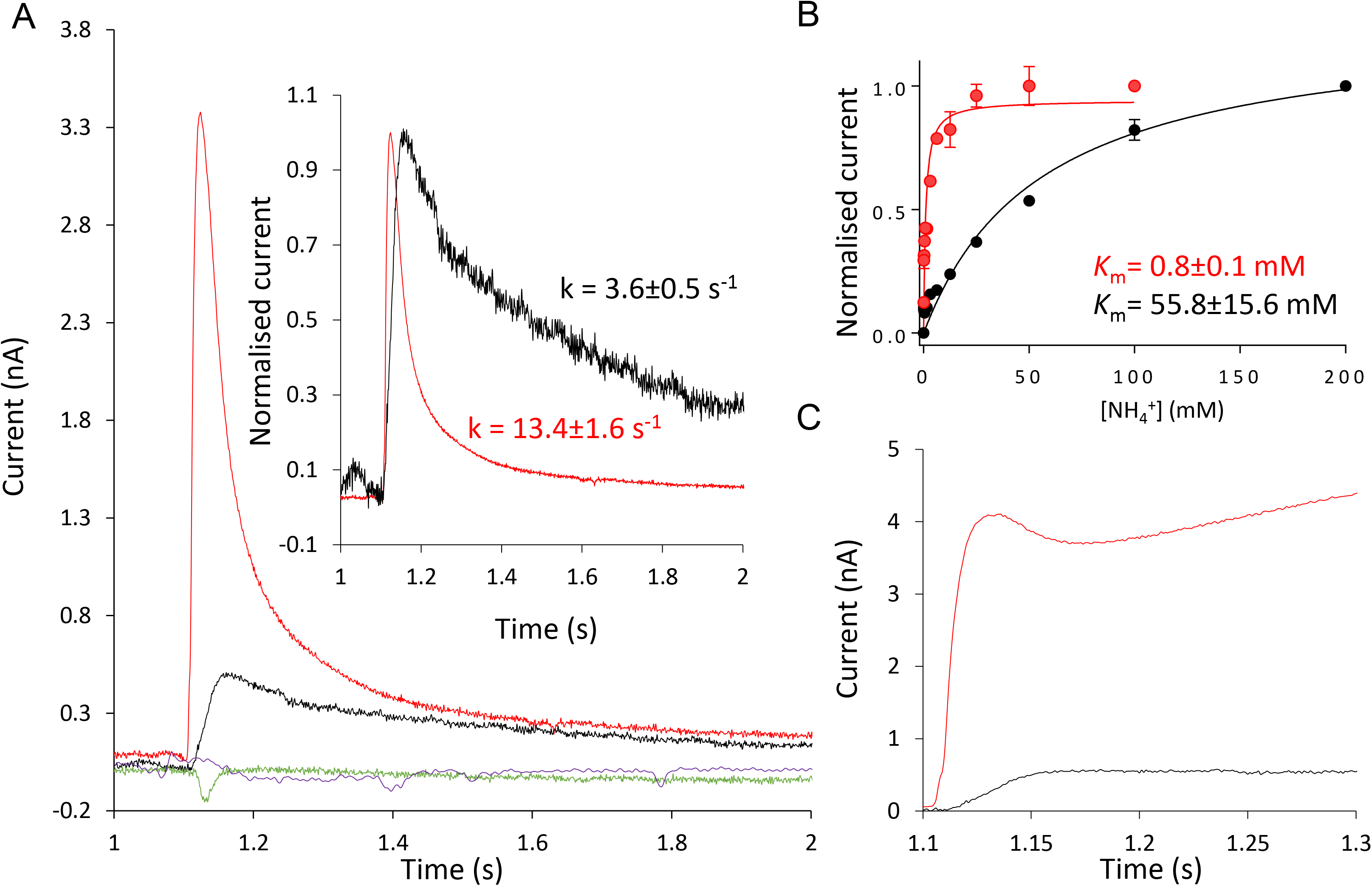
Specificity of AmtB activity. **(A)** Transient current measured on proteoliposomes containing AmtB at LPR 10 after a 100 mM substrate jump. Ammonium (red), methylammonium (black), potassium (green) or sodium (purple). ***Insert**:* Normalised current after a 100 mM substrate jump. Ammonium (red), methylammonium (black). **(B)** Substrate dependence (ammonium-red or methylammonium-black) of the maximum amplitude of the transient current **(C)** Reconstructed current using circuit analysis after a 100 mM jump of ammonium (red) or methylammonium (black).

To characterise the specificity of AmtB amongst monovalent cations, we measured the current after pulses of Na^+^ and K^+^. The ionic radii of Na^+^ (0.116 nm), and K^+^ (0.152 nm) are similar to the size of NH_4_^+^ ions (0.151 nm (28). In spite of this, 100 mM Na^+^ or K^+^ pulses did not trigger any charge displacement (Figure 3A). This shows that these ions neither interact with AmtB nor are they translocated through the protein. These experimental observations agree with previous free energy calculations, which have suggested a high energy barrier for the translocation of ions through AmtB due to the hydrophobicity of the pore (29–31).

Next, we investigated the specificity of AmtB for ammonium versus methylammonium (MeA) transport. MeA has been widely used to measure ammonium transport activity as the radioactive tracer ^[14]^C-MeA is commercially available. However, the suitability of MeA as an ammonium analogue to characterise the kinetics, specificity and energetics of Amt/Mep/Rh activity have been questioned (32, 33). Here, we show that a pulse of 100 mM MeA triggers a transient current of 0.50±0.02 nA, compared to 3.37±0.26 nA for ammonium, while the decay constant is 4 times lower (Figure 3A). The currents recorded by SSME are intrinsically transient; however, the transporter steady-state components can be reconstructed by circuit analysis (34). Steady-state transport in AmtB associated with 100 mM ammonium caused a current of ~4 nA, while for 100 mM MeA it was found to be ~0.5 nA. This shows that MeA is translocated through AmtB at a greatly reduced rate compared to ammonium (Figure 3C). Furthermore, kinetics analysis reveals a *K*_m_ 70 times higher (55.8mM) for MeA compared to ammonium, showing further that MeA is a poor substrate analogue for AmtB and not well suited to elucidate the mechanistic details of AmtB activity (Figure 3B).

### PG lipids are functionally required for AmtB transport activity

Given that PG has been shown to bind specifically to AmtB (23), we hypothesised that the lipid environment may also affect the protein’s activity, the ammonium-AmtB interaction and/or the translocation process. To test this question, we reconstituted AmtB into liposomes containing a mixture of phosphatidic acid (PA)/phosphatidylcholine (PC) at a weight ratio of 9:1 or ternary mixtures of PA/PC and PG. The ternary mixture was chosen such that the quantity of PG (16.5% wt/wt) matched the standard composition used for the previous experiments (*E. coli* polar lipids/PC 2/1 wt/wt, see table 1). IMAC analysis and DLS measurements showed that the size of the liposomes, the protein orientation inside the membrane and the quantity of protein inserted were equivalent for all lipid conditions (Figure S1). In the absence of PG (condition 2, table 1), a 100 mM ammonium pulse triggered a transient current of 0.42±0.04 nA, compared to 3.37±0.26 mA in the presence of PG (condition 1) (Figure 4A). Remarkably, in the presence of 16.5% PG in an otherwise pure PA/PC condition (condition 3) a current of 2.24±0.06 nA was measured, which is >5 times greater than in the absence of PG (condition 2) (Figure 4A). The decay constant and the *K*_m_smeasured in the presence of PG (conditions 1 and 3) were similar and remained within the experimental error. In contrast, in the absence of PG (condition 2), the decay rate and the *K*_m_ increased by 1. 6 and 7-fold, respectively, compared to conditions 1 and 3 (Figure 4A). All of these findings clearly show that PG lipids are required for the full transport activity of AmtB.

**Figure 4:**
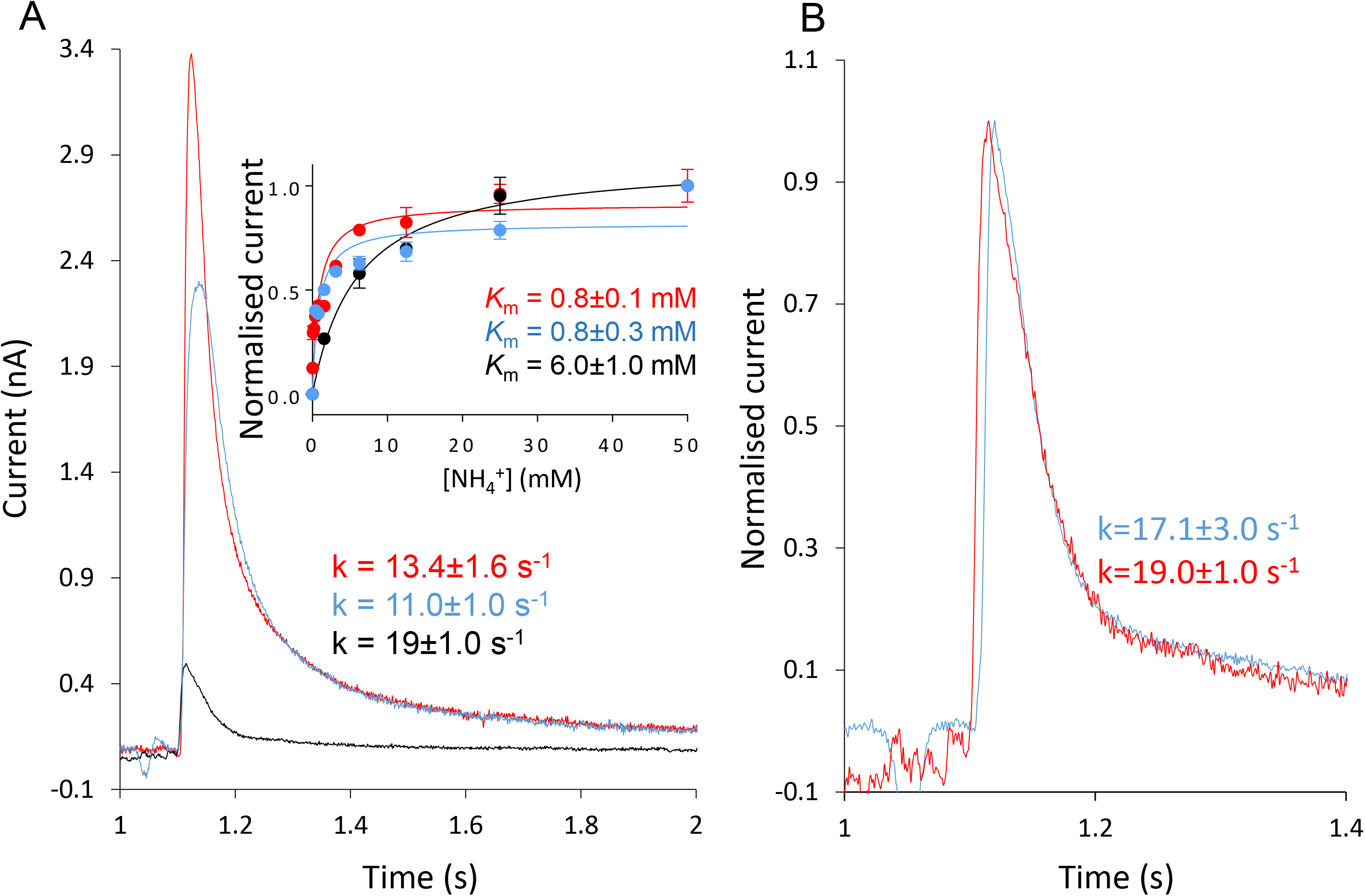
PG is required for the function of AmtB. **(A)** Transient current measured after a 100 mM ammonium jump in proteoliposomes containing the lipid conditions 2 (black), 3 (blue), 1 (red). ***Insert**:* Ammonium dependence of the maximum amplitude of the transient current for proteoliposomes containing the lipid conditions 2 (black), 3 (blue), 1 (red). **(B)** Normalised transient current measured in AmtB-containing proteoliposomes that do not contain PG (condition 2) at LPR10 (red) or 5 (blue).

**Table 1:** Lipid composition in liposomes and total PG content*

| Lipid conditions | Lipid content-ratio<br>(wt/wt) | % of PG |
| --- | --- | --- |
| 1 | *E. coli* polar/PC - 2/1 | 16.5 |
| 2 | PA/PC - 1/9 | 0 |
| 3 | PA-PC/PG - 5/1 | 16.5 |

* We reconstituted AmtB in liposomes containing a mixture of phosphatidic acid (PA)/phosphatidylcholine (PC) at a weight ratio of 9/1 (condition 2) or in PA/PC-containing liposomes from condition 2 but also containing PG at a weight ratio of 5/1 (condition 3). Condition 3 was chosen such that the quantity of PG (16.5% wt/wt) matched the standard composition used for the previous experiments (*E. coli* polar lipids/PC 2/1 wt/wt; condition 1).

To determine if AmtB completes a full transport cycle in the absence of PG, we reconstituted the protein in condition 2 (without PG, table 1) at varying LPRs of 50, 10 and 5 (wt/wt). Here, an ammonium pulse of 100 mM did not trigger a measurable transient current at LPR 50. However at LPR 10 and 5, the decay time was similar and within the experimental error (19.0±1.0 s^-1^ vs. 17.1±3.0 s^-1^ respectively; Figure 4B).

To ensure that AmtB was not misfolded in the liposomes under condition 2 (without PG), we solubilised the proteoliposomes in 2% DDM to extract AmtB and analysed the protein by SEC on a superdex 10/300 increase column. The SEC analysis showed that the protein elutes as a single monodisperse peak at the same retention time (11.6 ml) as prior to reconstitution under condition 2 (Figure 5A). We subsequently re-inserted AmtB solubilised from condition 2 (without PG) into proteoliposomes using condition 1 (with PG). An ammonium pulse of 100 mM in these proteoliposomes triggered a transient current of 3.18±0.09 nA with a decay constant of 11.5±2.4 s^-1^, and the kinetic analysis revealed a *K*_m_ of 1.3±0.3 mM (Figure 5B). These results show that AmtB reconstituted under condition 2 (without PG) regains the original activity parameters (maximum current intensity, decay time and *K*_m_) when re-reconstituted in liposomes under condition 1 (with PG). These data confirm correct folding of AmtB in the proteoliposomes without PG (condition 2). Taken together, these findings show that PG is essential for AmtB activity and furthermore indicate that in the absence of PG AmtB exhibits a defective transport cycle.

**Figure 5:**
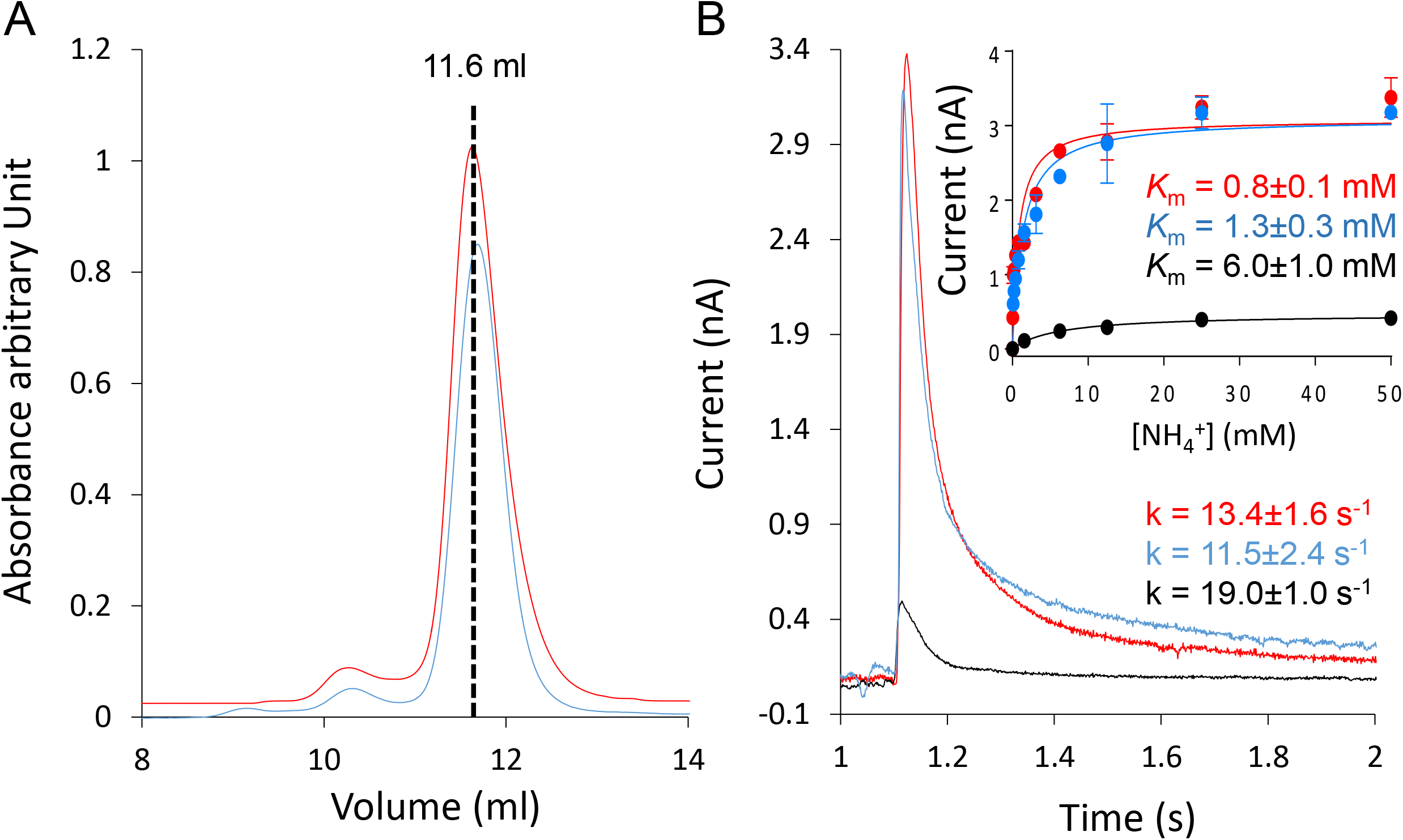
AmtB is correctly folded in absence of PG. **(A)** Gel filtration trace (Superdex 200 10/300 increase) of AmtB before (red) insertion in proteoliposomes under condition 2 and after (blue) solubilisation from proteoliposomes under lipid condition 2. **(B)** Transient current measured after a 100 mM ammonium jump in proteoliposomes under condition 2 (black), under condition 1 containing AmtB re-inserted after solubilisation from proteoliposomes under condition 2 (blue) and under condition 1 (red). ***Insert**:* Ammonium dependence (raw data) of the maximum amplitude of the transient current in proteoliposomes under condition 1 (red), 2 (black) or 4 (blue).

### Structural and dynamic investigation of PG interacting with AmtB

We next applied atomistic molecular dynamics (MD) simulations to study the interaction of PG lipids with AmtB in membranes on the molecular level. Figures 6A and S2A show the simulation systems containing an AmtB trimer embedded within PA/PC/PG and PA/PC mixed lipid membranes respectively. The colour maps on the right and bottom of figures 6A and S2 display the density of PA/PC and PG, respectively, each derived from 0.4 μs simulations.

**Figure 6:**
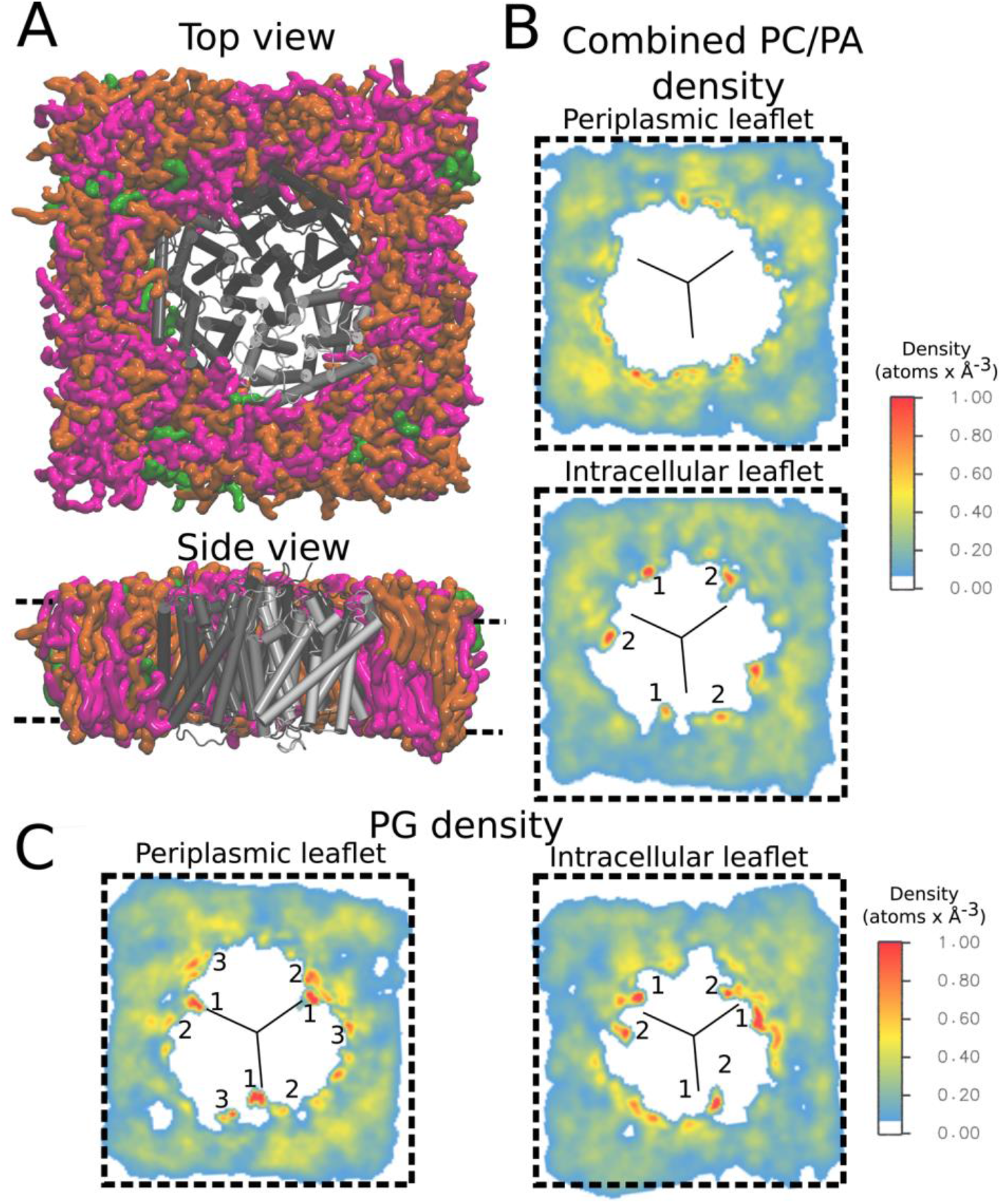
Trimeric AmtB in the PA/PC/PG system and lipid density plots. **(A)** Final frame of the simulation system, seen from the periplasm (top) and from the side (bottom). The protein is shown in grey, the PC lipid molecules in orange, PA in green, and PG in magenta. **(B)** Volumetric analysis of PC and PA combined average densities over the whole trajectory. **(C)** Volumetric analysis of PG average densities over the whole trajectory. The black lines in the density plots mark the approximate monomer interfaces. Specific binding sites are labelled in the density plots. A comparison of the 2-D density maps shows that PG tends to localize preferably close to the monomer interfaces.

Although specific AmtB interactions with PG had been crystallographically detected previously for the extracellular membrane leaflet (23), our simulations reveal additional PG binding sites on the intracellular side of the membrane (Figure 6B-C). PG lipids preferably occupy sites on the AmtB trimer located near the interfaces between the monomers, both within the periplasmic and intracellular membrane leaflets (Figure 6C). Specifically, we observe three high-density lipid interaction sites per subunit in the periplasmic leaflet, one at the monomer interface and two in its vicinity. Within the intracellular leaflet, about two high-density sites are seen per subunit, in which lipids interact with interfacial helices of AmtB in both cases (Figure 6C). By contrast, the density maps recorded for the PA/PC lipid mixture show no binding hotspots of AmtB for PC and PA within the periplasmic membrane leaflet, while some lipid accumulation is observed within the intracellular leaflet close to the AmtB PG binding sites 1 and 2 (Figure S2B). Close-up images highlighting the interaction between PG lipids and the AmtB subunits are shown in Figure 7. The binding sites observed in our MD simulations within the periplasmic leaflet are in good agreement with crystallographically defined sites (23) (Figure 7A), while the new intracellular interaction sites also locate to the interface region between the AmtB subunits (Figure 7B).

**Figure 7:**
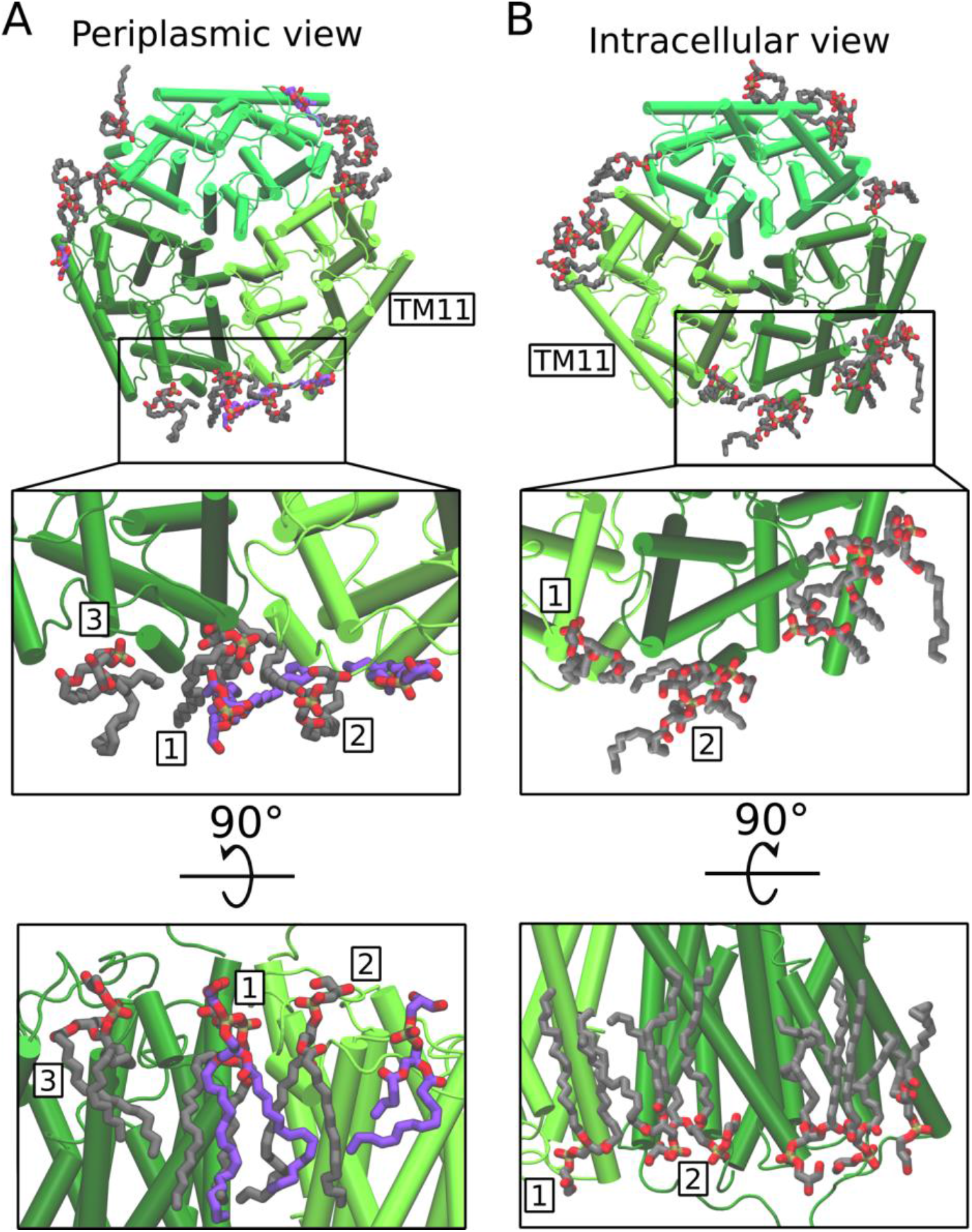
PG binding sites on AmtB. The images highlight PG binding locations, taken from a representative simulation frame of the PA/PC/PG mixture. **(A)** PG binding sites located in the periplasmic membrane leaflet. The simulated PG molecules bound to AmtB are shown in grey, while the crystallographically resolved bound lipids (PDB ID: 4NH2) are in violet. Good agreement between the experimental and the simulation sites is observed, especially for binding site 1. **(B)** PG binding sites in the intracellular leaflet. The binding sites are labelled according to the volumetric maps shown in Figure 6. Generally, PG tends to bind close to the monomer interface.

The most conspicuous conformational change we find to be induced by PG binding in AmtB on the time scale of our simulations is the formation of a short helix within an external loop region (residue 80 to 85) following binding of the charged PG headgroup (Figure S3). Additionally, the loops 157-166 and 331-348 display an enhanced dynamic behaviour in PG-containing lipids when compared to PA/PC (Figure S3A). Notably, loop 149-163 includes the residue D160, which has experimentally been shown to be essential for the transport activity of AmtB (29, 30, 35, 36) (Figure S3B). Although the detailed mechanism of ammonium transport in AmtB is still unclear, this finding might link PG binding to AmtB function.

## Discussion

A plethora of functional studies aimed at elucidating the mechanism of ammonium transport by Amt proteins has led to considerable controversy due to the lack of an *in vitro* assay characterizing the activity of Amt proteins using ammonium as substrate (1, 9). The elegant work of Wacker *et al.*, (2014) showed that two ammonium transporters from *Archaeoglobus fulgidus* activity are electrogenic (37). The work we present here shows that the activity of the archetypal Amt/Mep/Rh protein, AmtB, is also associated with charge translocation, which suggests that this may be a general feature for these proteins. However, we also demonstrate that contrary to *A. fulgitus* Amt1, AmtB highly discriminates between MeA and ammonium as substrate. While in AfAmt1 MeA triggers a transient current amounting to 87% of the current elicited by ammonium (37), the current induced by MeA in AmtB is less than 15% of that observed for ammonium. The structural basis of this discrepancy is not clear since the three principal conserved features, namely the S1 binding site, the “phenylalanine gate” and the pore twin-His motif are structurally similar amongst both proteins. This functional difference in between eubacterial and archaeal Amts raises important questions about the universality of the transport mechanism in microbial ammonium transporters. This is particularly relevant in the case of the Mep-2 like protein that has been assigned a sensor role in filamentous development, often related to the virulence of pathogenic fungi (13). There are two hypotheses concerning the molecular mechanism of Mep2-mediated signaling. The first is that Mep2 is a sensor interacting with signaling partners leading to filamentation induction (38, 39) and the other that the transport mechanism of Mep2-like proteins may differ in Mep proteins (40–42). Current evidence favors the second hypothesis given that signalling efficiency is closely linked to transport efficiency, however further studies are needed to elucidate the exact mechanism, which may provide important information for the design of novel antifungal therapies.

It is well established that lipids can impact membrane protein structure and function through bulk membrane effects, by direct but transient annular interactions with the bilayer-exposed surface of protein, or by specific lipid binding to protein sites (for review see Ref. (43)). Altogether eight molecules of PG have been resolved in a recent crystal structure of AmtB, interacting at specific binding sites within the extracellular membrane leaflet (23). Our molecular dynamics simulations identified further interactions of PG molecules in the inner leaflet of the membrane. However so far, no functional relationships have been reported to link PG binding with the activity of AmtB.

Here, we show that in the absence of PG, AmtB is non-functional as a transporter and unable to complete the full translocation cycle. Our experiments and simulations do not indicate substantial conformational changes in the S1 periplasmic binding site or in the pore, suggesting that the molecular basis of the PG effect on AmtB activity could involve novel mechanistic sites. In this context, it is interesting to compare our findings with the lactose permease LacY in *E. coli*, the most extensively studied secondary transporter with regard to lipid-protein interactions. In the absence of phosphatidylcholine (PC) and/or phosphatidyethanolamine (PE), LacY is unable to support active transport of the substrate into the cell, although the binding of the substrate to the protein is unaffected (44). This is similar to the observations we made for AmtB in the absence of PG lipids. LacY is known to undergo drastic topological rearrangements, which may explain the effect of PC/PE on LacY activity (45). In the case of AmtB, no topological rearrangements were observed upon PG binding and we have shown that, in the absence of PG, AmtB was folded correctly in the proteoliposomes, such that a major change in the AmtB topology is unlikely to explain the functional role of PG. We do, however, observe changes in the dynamics of AmtB subunits, which locate particularly to loop regions at the periplasmic face of the protein. These regions include functionally important residues, for instance D160 (29, 30, 35, 36). Once a more detailed picture of the overall transport mechanism of AmtB has been obtained, the role of these effects may emerge in greater clarity.

Furthermore, several lines of evidence point towards functional cooperativity between the three subunits in the Amt/Mep transporters. Firstly, in *S. cerevisiae*, it has been demonstrated that expression of a non-functional Mep1 protein inhibits the transport activity of Mep2 and Mep3, indicating cross-talk between different Mep transporters (46), and similar observations have been reported for ammonium transporters of *Aspergillus nidulans* (47). Secondly, coexpressed non-functional monomers cross-inhibit transport in plant Amts, and a genetic screen has identified several mutations at the subunit interface of the *Arabidopsis thaliana* Amt1;2 transporter which inactivate the translocation activity (8, 48). Thirdly, extensive site-directed mutagenesis of the C-terminal tail of *Ec*AmtB has led to the hypothesis that the three subunits function in a cooperative manner (49). All of these findings indicate functional coupling between the adjacent subunits. Previous structural data and our MD simulations show that PG molecules bind mainly to sites at the vicinity of the subunit interfaces. It is then very attractive to hypothesise that PG molecules act as wedges, which mediate the functional interaction between the subunits. In line with this hypothesis, a model in which various AmtB conformations may be favoured upon specific lipid binding has been proposed (24). Finally, it has been shown recently that other lipids, including PE and cardiolipin, can bind AmtB allosterically, indicating that transporters may recruit their own micro-lipidic environment (50). Whether these binding events are important to modulate the activity of AmtB remains a question to be addressed in future studies.

## Methods

### Protein purification

AmtB(His_6_), cloned into the pET22b vector, was overproduced and purified as described (9) previously except that 0.03% of n-Dodecyl-β-D-Maltoside (DDM) was used instead of 0.09% *N*,*N*-dimethyldodecylamine-*N*-oxide (LDAO) in the final Size Exclusion Chromatography buffer (Tris/HCl 50mM, pH7.8, NaCl 100mM, 0.03% DDM) (51). AmtB was kept in the SEC buffer at 4°C for subsequent characterisation/ insertion into proteoliposomes.

### Reconstitution in liposomes

All lipids (Avanti polar lipids) were dried out under nitrogen flow and re-suspended at 5 mg/ml in non-activating (NA) buffer (Table 2). The multilamellar liposomes were subsequently extruded 13 times using the Mini-extruder (Avanti polar lipids) mounted with a 0.1 μm filter pore. To facilitate the insertion of AmtB in liposomes, 1 μl Triton X-100 at 25% was sequentially added to 500 μl of liposomes and the absorbance at 400, 500, 550 and 600 nm was measured in order to determine the R_sat_ and R_sol_ constants. The liposomes were incubated for 5 minutes at room temperature. AmtB, stabilised in 0.03% DDM, was added at a lipid:protein ratio of 5:1, 10:1 or 50:1 (wt/wt) and the mixture left for 30 minutes at room temperature. Three subsequent incubations with pre-washed SM-2 Biobeads (Biorad) at a beads:detergent ratio (wt/wt) of 20 were carried out to ensure detergent removal and AmtB insertion. The average diameter of the liposomes/ proteoliposomes was determined by dynamic light scattering (DLS) using a Zetasizer Nano ZS (Malvern instruments). Proteoliposomes were aliquoted in 100 μl and frozen at −80°C.

**Table 2:** Buffer composition of activated and non-activated solution for the SSME experiment*

|  |  |
| --- | --- |
| Non-activated solution | 100 mM potassium phosphate pH 7<br>300 mM KCl |
| Activated solution | 100 mM potassium phosphate pH 7<br>300-X mM KCl<br>X mM of substrate (NH<sub>4</sub><sup>+</sup>, MeA, Na<sup>+</sup>, K<sup>+</sup>) |

*When Na^+^ was used as a substrate, all of the buffers were prepared in Na-phosphate/NaCl

### AmtB orientation

160 μl of proteoliposomes (5 mg/ml) were treated with or without 2% DDM were incubated with 100 μl of Ni-affinity resin (Ni-Sepharose High Performance, GE healthcare) at 4°C for 1 hour. The supernatant was collected and the resin was washed 4 times using 50 μl of NA buffer (Table 2). The proteoliposomes were eluted in NA buffer containing 500 mM imidazole. 15 μl of each fraction were mixed with 5 μl of loading blue buffer and analysed by SDS-PAGE.

### Sensor preparation for SSME measurement

3-mm gold coated sensors were prepared as previously described (52). Briefly, 50 μl of a 0.5 mM octadecanethiol prepared in iso-propanol was used to coat a thiol layer on the gold surface of the sensor during 30 minutes. The sensors were rinsed with iso-propanol and deionised water, dried and diphytanoyl-sn-glycerol-3-phosphocholin solution was dropped onto the surface. 100 μl of non-activating (NA) buffer (Table 2) was immediately added to the sensor to form the solid supported membrane (SSM). Proteoliposomes/ empty liposomes were defrosted and sonicated in a sonication bath (U300H Ultrawave Precision ultrasonic cleaning) at 35 W during 1 minute, diluted 10 times in NA buffer (Table 2) and 10 μl was added at the surface of the SSM on the sensor. After centrifugation, the sensors were stored at 4°C for a maximum of 48 hrs prior to electrophysiogical measurements.

### SSME measurement

The measurements were performed using a SURFE^2^R N1 machine (Nanion Technology GmBH) using default parameters (26). The quality of the sensor was assessed prior to any measurement by determining its capacitance (value should be between 15 - 30 nF) and conductance (value should be below 5 nS). For the measurement, a single solution exchange was used that consisted of 3 phases of 1s each during which non-activating (NA), activating **(A)** and NA buffer were sequentially injected on the sensor over constant osmolality (Table 2) at a flow rate of 200μL/s. The sample rate was set to 1000 Hz, and all the currents in the figures are presented at this rate, without filtering. The currents were amplified with a gain set to 10^9^ V/A.

Wherever stated, the raw transient curves were normalised against the maximum current and for the kinetics against the maximum current recorded after a substrate pulse of 100 mM.

The measurements were done at least two times on each sensor, and on 6 sensors prepared from 2 independent batch of AmtB purification. The decay constant was fitted using Origin (OriginLab Corporation) and the kinetics analysis was done using Prism7 (GraphPad). The current reconstruction was done as previously described (52).

### Molecular Dynamics Simulations

The AmtB crystal structure at 1.35 Å resolution (pdb ID: 1U7G) (53) was used for all of our molecular dynamics simulations. The CHARMM_GUI web server (54, 55) was applied to revert the mutations inserted during the crystallization process back to the wild type form. The protein termini were capped with acetyl and N-methyl moieties for the N-terminus and C-terminus, respectively. The protein was then inserted into a 13×13 nm membrane patch (54).

Three different membrane compositions were used: one containing PA and PC lipids (PA/PC 1:9), and two containing PA, PC, and PG lipids (PA/PC/PG 1:9:2 and 1:9:10). K^+^ and Cl^-^ ions were added to neutralise the system and to obtain a bulk ionic concentration of 150 mM.

The CHARMM36 force field was used for the protein, lipids and ions (56, 57). The water molecules were modelled with the TIP3P water model (58). Water bonds and distances were constrained by the Settle method (59), and all other bonds by the LINCS method (60). After a steepest descent minimisation, the system was equilibrated by six consecutive equilibration steps with position restraints on heavy atoms of 1000 kJ/mol nm^2^. The first three equilibration steps were carried out in a NVT ensemble using a Berendsen thermostat (61) to keep the temperature at 310K. The subsequent steps were conducted under a NPT ensemble, switching on a Berendsen barostat (61) with isotropic coupling, to keep the pressure at 1 bar. Production molecular dynamics simulations were carried out using a Nose-Hoover thermostat (62) with a time constant of 0.2 ps, and a Parrinello-Rahman barostat (63, 64) with semiisotropic pressure coupling. An integration time step of 2 fs was used throughout the simulations. The VMD software (65) was used for the visualisation of the trajectories and the generation of all structural images. We used the VMD VolMap plugin for the generation of the volumetric density maps.

## Author contributions

G.D. Mirandela, G. Tamburrino, U. Zachariae, A. Javelle designed and performed the research and analysed the data. A, Javelle, U. Zachariae and P.A. Hoskisson wrote the paper.

## Acknowledgement

GDM was supported by a PhD studentship from the University of Strathclyde. AJ was supported by a Chancellor’s Fellowship from the University of Strathclyde, GT and UZ acknowledge funding from the Scottish Universities’ Physics Alliance (SUPA). AJ acknowledges the support of Tenovus Scotland (project S17-07) and PAH would like to acknowledge the support of the Natural Environment Research Council (Grant: NE/M001415/1). Thanks to Dr. Andre Bazzone (Nanion Technologies, GmbH) for helpful advice with SSME measurement. Special thanks to Prof. Iain Hunter (Strathclyde Institute of Pharmacy and Biomedical Sciences) for invaluable discussions and help during this project.

